# Ligation of random oligomers leads to emergence of autocatalytic sequence network

**DOI:** 10.1101/2020.08.18.253963

**Authors:** Patrick W. Kudella, Alexei V. Tkachenko, Sergei Maslov, Dieter Braun

## Abstract

The emergence of longer information-carrying and functional nucleotide polymers from random short strands was a major stepping stone at the dawn of life. But the formation of those polymers under temperature oscillation required some form of selection. A plausible mechanism is template-based ligation where theoretical work already suggested a reduction in information entropy.

Here, we show how nontrivial sequence patterns emerge in a system of random 12mer DNA sequences subject to enzyme-based templated ligation reaction and temperature cycling. The strands acted both as a template and substrates of the reaction and thereby formed longer oligomers. The selection for templating sequences leads to the development of a multiscale ligation landscape. A position-dependent sequence pattern emerged with a segregation into mutually complementary pools of A-rich and T-rich sequences. Even without selection for function, the base pairing of DNA with ligation showed a dynamics resembling Darwinian evolution.

## BACKGROUND

One of the dominant hypotheses to explain the origin of life^1–3^ is the concept of RNA world. It is built on the fact that catalytically active RNA molecules can enzymatically promote their own replication^4–6^ via active sites in their three dimensional structures^7–9^. These so-called ribozymes have a minimal length of 30 to 41 bp^9,10^ and, thus, a sequence space of more than 4^30^≈10^18^. The subset of functional, catalytically active sequences in this vast sequence space is vanishingly small^11^ making spontaneous assembly of ribozymes from monomers or oligomers all but impossible. Therefore, prebiotic evolution has likely provided some form of selection guiding single nucleotides to form functional sequences and thereby lowering the sequence entropy of this system.

The problem of non-enzymatic formation of single base nucleotides and short oligomers in settings reminiscent of the primordial soup has been studied before^12–16^. However, the continuation of this evolutionary path towards early replication networks would require a pre-selection mechanism of oligonucleotides (as shown in Fig. 1a), lowering the information entropy of the resulting sequence pool^17–20^. In principle, such selection modes include optimization for information storage, local oligomer enrichment e.g. in hydrogels or in catalytically functional sites.

**Fig. 1.**
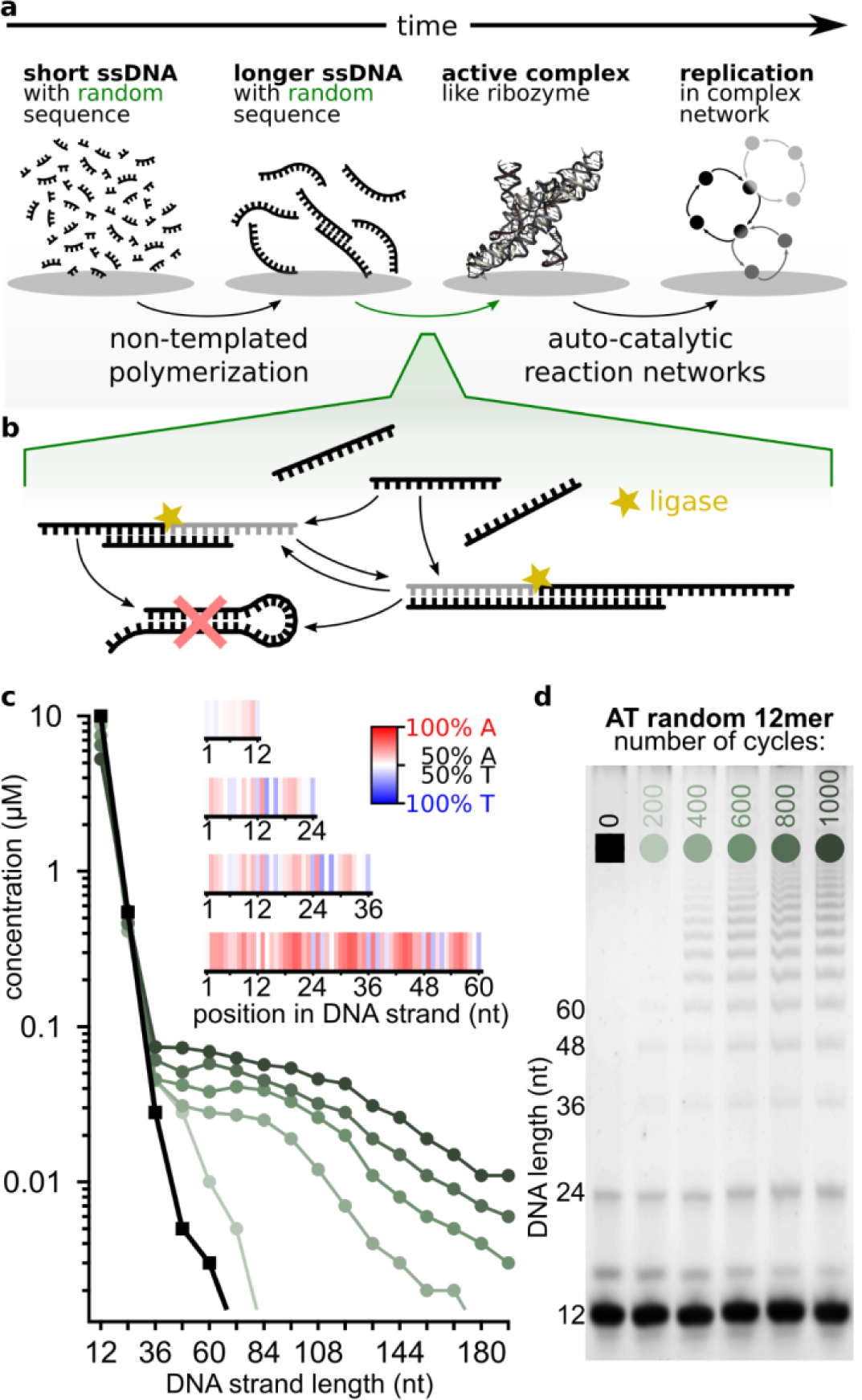
Autocatalytic templated ligation of DNA 12mers. **a** Before cells evolved, the first ribozymes were thought to perform basic cell functions. In the exponentially vast sequence space, spontaneous emergence of a functional ribozyme is highly unlikely, therefore pre-selection mechanisms were likely necessary. **b** In our experiment, DNA strands hybridize at low temperatures to form 3D complexes which can be ligated and preserved in the high temperature dissociation steps. The system self-selects for sequences with specific ligation site motifs as well as for strands that continue acting as templates. Hairpin sequences are therefore suppressed. **c** Concentration analysis shows progressively longer strands emerging after multiple temperature cycles. The inset (A-red, T-blue) shows that while 12mers (88009 strands) have essentially random sequences (white), various sequence patterns emerge in longer strands (60mers, 235913 strands analyzed). **d** Samples subjected to different number (0-1000) of temperature cycles between 75 °C and 33 °C. Concentration quantification is done on PAGE with SYBR post-stained DNA.

An important aspect of a selection mechanism is its non-equilibrium driving force. Today’s highly evolved cells function through multistep and multicomponent metabolic pathways like glycolysis in the Warburg effect^21^ or by specialized enzymes like ATP synthase which provide energy-rich adenosine triphosphate (ATP)^22^. In contrast, it is widely assumed^3,4,23–26^ that selection mechanisms for molecular evolution at the dawn of life must have been much simpler, e.g. mediated by random binding between biomolecules subject to non-equilibrium driving forces such as fluid flow and cyclic changes in temperature.

Here, we explored the possibility of a significant reduction of sequence entropy driven by templated ligation^17^ and mediated by Watson-Crick base pairing^27^. Starting from a random pool of oligonucleotides we observed a gradual formation of longer chains showing reproducible sequence landscape inhibiting self-folding and promoting templated ligation. Here we argue, that base pairing combined with ligation chemistry, can trigger processes that have many features of the Darwinian evolution.

As a model oligomer we decided to use DNA instead of RNA since the focus of our study is on base pairing which is very similar for both^28^. We start our experiments with a random pool of 12mers formed of bases A (adenosine) and T (thymine). This binary code facilitates binding between molecules and allows us to sample the whole sequence space in microliter volumes (2^12^ << 10 µM * 20 µl = 10^14^).

Formation of progressively longer oligomers from shorter ones requires ligation reactions, a method commonly employed in hairpin-mediated RNA and DNA replication^29,30^. At the origin of life, this might have been achieved by activated oligomers^31,32^ or activation agents^33–35^. Our study is focused on inherent properties of self-assembly by base pairing in random pools of oligomers and not on chemical mechanisms of ligation. Hence, we decided to use TAQ DNA ligase - an evolved enzyme for templated ligation of DNA^19^. This allowed for fast turnovers of ligation and enabled the observation of sequence dynamics.

## RESULTS

To test templated elongation of polymers in pools of random sequence oligomers, we prepared a 10 µM solution of 12mer DNA strands composed of nucleotides A and T (sequence space: 4096) and subjected it to temperature cycling, similar to reference^19^ with 20 s at denaturation temperature of 75 °C and 120 s at ligation temperature of 33 °C. Temperatures were selected according to the melting dynamics of the DNA pool; the time steps were prolonged relative to Toyabe and Braun (SI section 5.3) because of a greater sequence space. The sample was split into multiple tubes and exposed to 200, 400, 600, 800, 1000 temperature cycles, with one tube kept at 4 °C for reference.

To study the length distributions in our samples we used polyacrylamide gel electrophoresis (PAGE, Fig. 1d). The first lane is the reference sequence not exposed to temperature cycling, where small amounts of impurities are visible at short lengths (SI Section 3.1). The latter lanes show the temperature-cycled samples. As the number of cycles increases, progressively longer strands in multiples of 12 emerge, as the original pool only consisted of 12mers. Fig. 1c shows the concentration quantification of each lane (compare SI section 3). For higher cycle counts the total amount of products increases and the concentration as a function of length decreases slower. The behavior of this system is dependent on the time and temperature for both steps in the temperature cycle, the monomer-pool concentration and the sequence space of the pool (SI section 5).

An important property of the initial monomer-pool is its sequence content. Although for pools with lower sequence complexity it is possible to show different strand compositions using PAGE^36,37^, a large size of our “monomer” (2^12^=4096) and 24mer product pools (sequence space: 2^24^≈16.8×10^6^) excludes this approach. Thus, we analyzed our final products by Next Generation Sequencing (NGS) to get insights into product strand compositions.

Plotting the probability of finding a base at a certain position (Fig. 1c inset) revealed no distinct pattern in 12mers other than a slight bias towards As. However, longer chains starting with 24mers developed a strikingly inhomogeneous sequence pattern: bases around ligation sites show a distinct AT-alternating pattern, while regions in the middle of individual 12mers are preferentially enriched with As.

The information entropy of longer chains is expected to be smaller than the entropy of a random sequence strand of the same length, if some sort of selection mechanism is involved^17^. We analyzed the entropy reduction for different lengths of products (Fig. 2a) as well as the positional dependence of the single base entropy for 60mer products (Fig. 2b). The relative entropy reduction is similar to one used in Derr *et. al*^38^ where 1 describes a completely random ensemble and 0 an ensemble of only one sequence. Entropy reduction was observed in all analyzed product lengths with a greater reduction observed for longer oligomer lengths. The entropy of each 12mer subsequence was also found to be significantly lower than that of random 12mers (Fig. 2b, black line). The central subsequence had the lowest entropy while 12mers located at both ends of chains had relatively higher entropies. This behavior was also observed as a function of nucleotide position within a 12mer suggesting a multi-scale pattern of entropy reduction.

**Fig. 2.**
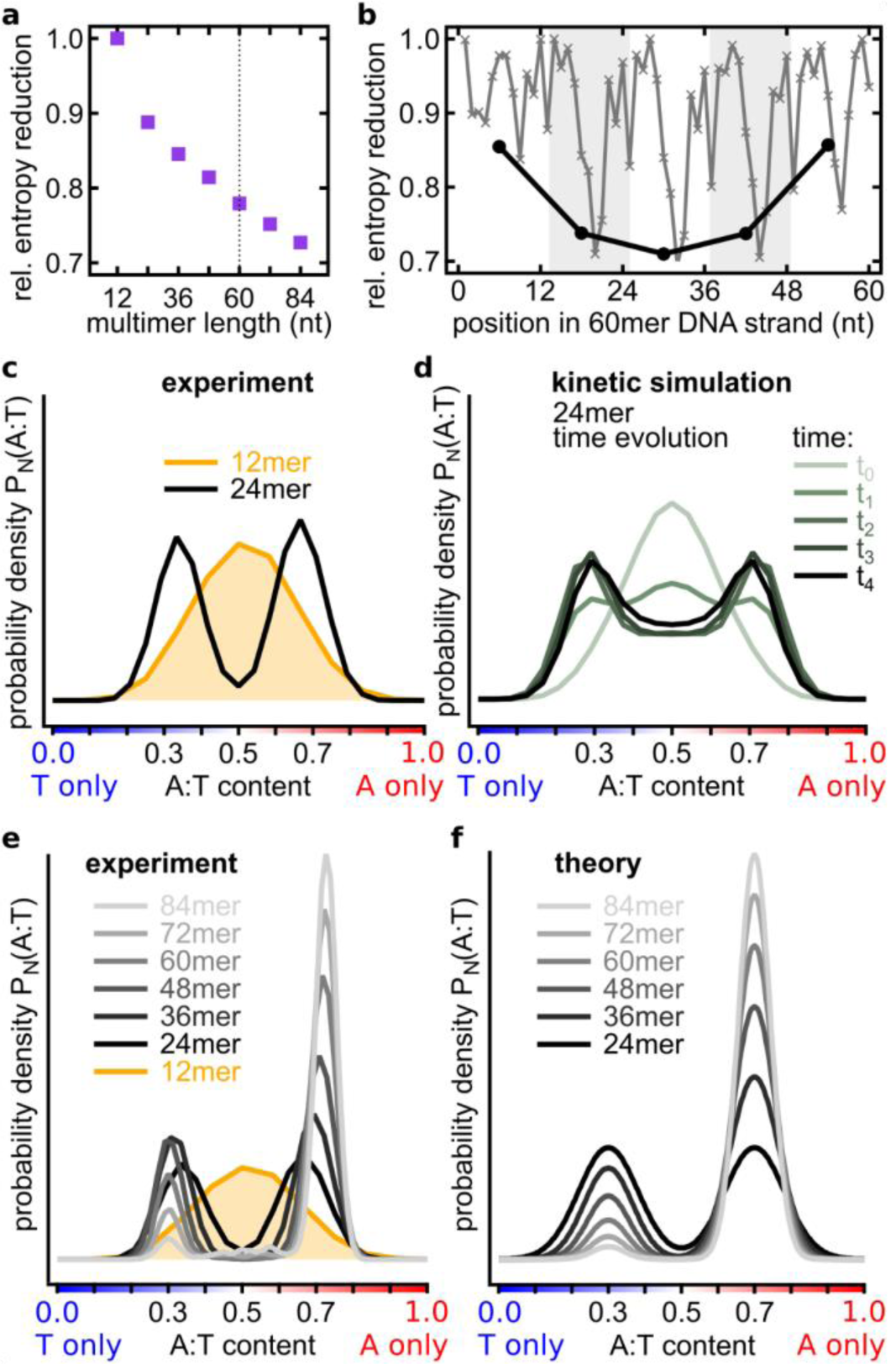
Hairpin formation amplifies selection into A-rich and T-rich sequences. **a** Relative entropy reduction as a function of multimer product length: 1 – a random pool and 0 – a unique sequence. **b** Relative entropy reduction of 60mer products. Black: Entropy reduction of 12 nt subsequences compared to a random sequence strand of the same length. Grey: Entropy reduction at each nucleotide position showing positional dependence. **c** A gradual development of the bimodal distribution of A:T ratio in chains of different lengths. While the A:T ratio in 12mers has a single-peaked nearly binomial distribution, 24mers already have a clearly bimodal distribution peaked at 65:35 % (A-type strands) and 35:65 % (T-type strands) A:T ratios. **d** Emergence of a bimodal distribution in a kinetic model of templated ligation. Sequences with nearly balanced A:T ratios are prone to formation of hairpins. In the model these hairpins prevent strands from acting as templates and substrates for ligation reactions thereby suppressing the central part of the distribution. **e** A:T ratio distributions in strands of different length. As length increases A-type strands become progressively more abundant in comparison to T-type strands. **f** A:T ratio distributions in a phenomenological model taking into account a slight AT-bias in the initial 12mer pool resemble experimentally measured ones (panel e).

In the initial pool of random 12mers the A-to-T ratio distribution is shaped binomially, as expected for a random distribution. However, it dramatically shifted for 24mer products of ligation: a bimodal distribution of about 65:35 % A:T (A-type) as well as the inverse, 35:65 % A:T (T-type) was observed with 24mer products (Fig. 2c). DNA strands composed of only two complementary bases are more prone to formation of single-strand secondary structures like hairpins than DNAs composed of all four bases. In our templated ligation reaction, we expected that hairpin-sequences are not elongated and also not used as template-strands because they form catalytically passive Watson-Crick-base-paired configuration. A bimodal AT-ratio distribution (Fig. 2d) also emerged in a kinetic computational model in which a pool of random 12mers was seeded with a small initial amount of random sequence 24mers. 24mers that formed hairpins could not act as templates and were therefore less likely to be reproduced (see SI for details of this model, section 18.2).

For longer products the bimodal distribution got sharper and centered at approximately 70:30 % A:T and 30:70 % A:T (Fig. 2e). To compare the distributions of different lengths we computed probability density functions (PDF) of A:T fractions. Each distribution is the sum (integral) over all probabilities *P*_*N*_ to find a certain A:T-fraction d_*A*:*T*_ in chains of length *N*:

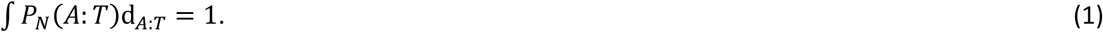

The main difference of longer oligomers was a rapid increase of the ratio between the number of A-type and T-type sequences. As oligomers get longer the effect becomes more pronounced. This might be a result of a small bias in the initial pool which has slightly more monomers of A-type than T-type (SI section 9.1).

As predicted theoretically^39^, the eventual length distribution is approximately exponential. A small A-T bias leads to the respective average chain lengths, 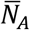 and 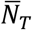, to be somewhat different for the two subpopulations. As a result, the bias gets strongly amplified with increasing chain length:

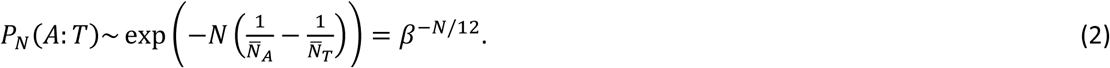

A simple phenomenological model can successfully capture the major features of the observed A:T PDFs for multiple chain lengths. Specifically, we assume both A-type and T-type sub-populations -to maximize the sequence entropy, subject to the constraint that the average A:T content is shifted from the midpoint (50:50 % composition), by values ±*x*_0_, respectively. This model presented in SI section 18.1, results in a distribution that strongly resembles experimental data, as shown in Fig. 2e-f A:T profiles for all chain length are fully parameterized by only two fitting parameters: *β* = 0.785, and *x*_0_ = 0.2.

The proposed mechanism of selection of A-type and T-type subpopulations due to hairpin suppression is further supported by direct sequence analysis. Fig. 3b shows PDFs of the longest sequence motifs that would allow hairpin-formation, across the entire pool of sequences of given lengths. While the overall chain length increased by a factor of seven (12 to 84 nt), the most likely hairpin length only grew by a factor of 1.89 (3.7 to 7 nt) (Fig. 3b). The observed relationship between the strand length *N* and the most likely hairpin length *l*_0_ can be successfully described by a simple relationship obtained within the above described maximum-entropy model. Specifically, for a random sequence with bias parameter *p* = 0.5 + *x*_0_, one expects *N* to be related to *l*_0_ as follows (as in Fig. 2f):

**Fig. 3.**
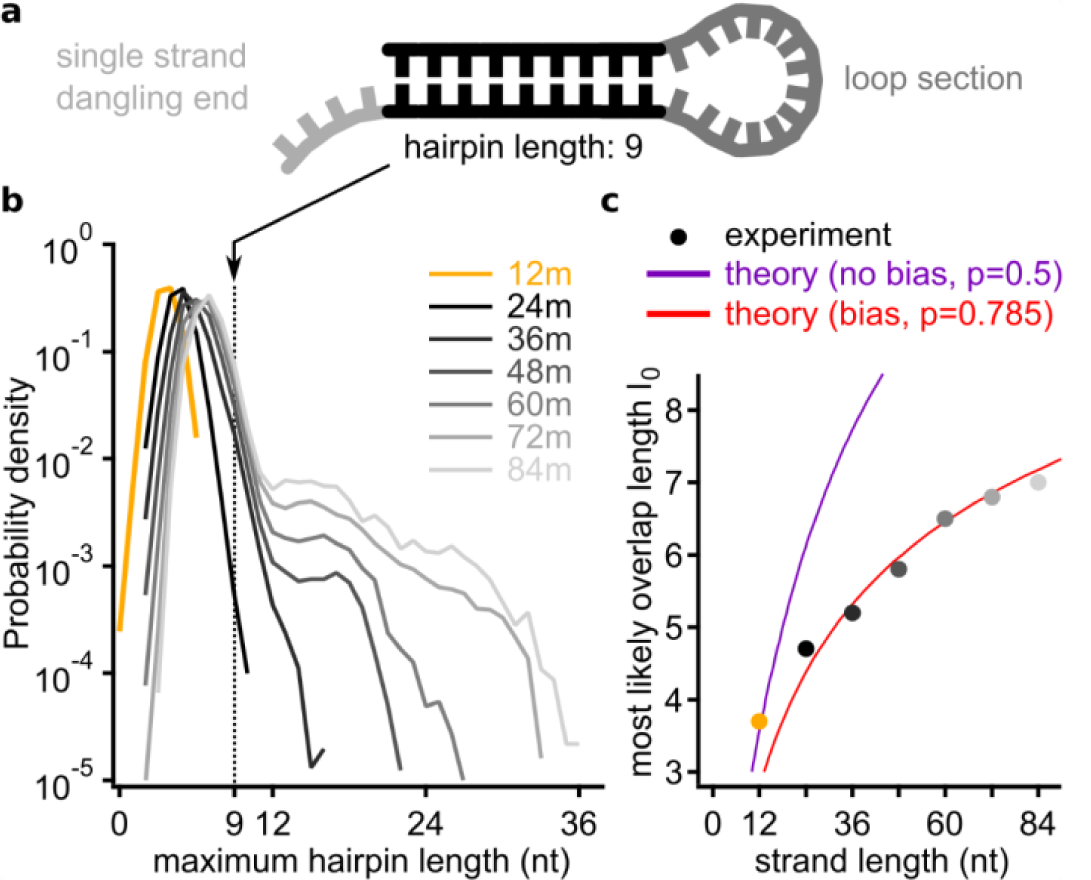
Large scale entropy reduction and sequence correlation per strand. **a** Sketch of a single strand DNA secondary structure folding on itself, called hairpin. The double stranded part is very similar to a standard duplex DNA. **b** Comparing the PDFs of the maximum hairpin length for all strands reveals a group of peaks at around 4 to 7 nt, increasing with the DNA length. Starting for 48mers, there is a tail visible: these self-similar strands are more abundant, the longer the product grows (compare A:T fraction close to p=0.5 in Fig. 2c). **c** The peak-positions as function of the product length follow equation (3). The unbiased 12mers are on the curve with coefficient p=0.5, whereas the products starting from 36mers lay on the curve with p=0.785. The bias parameter p is derived from the PDFs in Fig. 2d and describes the A:T-ratio in the strand.

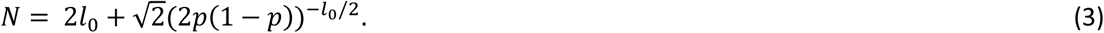

As one can see in Fig. 3c, this result is in an excellent agreement with experimental data for all the long chains, assuming *p*=0.785. This A:T ratio is indeed comparable to the one observed in the A-type subpopulation. On the other hand, the maximum probability length of the longest hairpin for 12mers is consistent with an unbiased composition, *p*=0.5.

While hairpin formation inhibits the self-reproduction based on template-based ligation, Fig. 3b reveals another dramatic feature: a small fraction of chains does feature very long hairpin-forming motifs (seen as shoulders in the distribution function). This effect also reveals itself as small peaks on the 84mer curve in Fig. 2e. Those peaks around A:T ratio 0.4, 0.5 and 0.6. stem from subpopulations that have multiple A-types as well as multiple T-type subsequences (see SI section 12) and are prone to hairpin formation.

The mechanism of formation of these self-binding sequences may involve recombination of shorter A-type and T-type chains, or self-elongation of shorter hairpins. In either case, the harpin sequence cannot efficiently reproduce by means of template ligation. However, the reminder of the pool would keep producing them as byproduct. Ironically, for the templated ligation reaction this is a possible failure mode, but those long hairpins may play a key role in the context of origin of life, as precursors of functional motifs. For instance, work by Bartel and Szostak^11,40^ identifies RNA self-binding as crucial for the direct search of ribozymes – those molecules need to fold into non-trivial secondary structures to gain their catalytic function.

The separation into A-type and T-type subpopulations only accounts for a small part of the sequence entropy reduction. The emerging ligation landscape in the sequence space is far richer.

Sequence analysis of the junctions in-between original 12mer revealed additional information about that landscape, already hinted by patterns seen in Fig. 1b. We characterize pairs of junction-forming sequences with their Z-scores, i.e. probability of their occurrence scaled with its expected value and divided by the standard deviation calculated in the random binding model (see SI section 14).

Fig. 4a shows Z-score heatmaps for junctions within A-type (left panel) and T-type (right panel) subpopulations. More specifically, we show sequences left (row) and right (column) of the junction between the 4th to the 5th 12mers in the respective 72mer. These heatmaps reveal a complex landscape of over- and under-represented junction motifs shown respectively in dark-teal and dark-ocher colors. Emergence of such complex landscape has been theoretically predicted in Ref. ^17^ Landscape peaks include repeating A-T motif of alternating bases crossing the ligation site (dark-teal peak near the center of each of both heatmaps). Relatively rare motifs (valleys) correspond to poly-A and poly-T sequences extending across the junction (dark-ocher areas). One exception to this rule is a relatively abundant poly-A motif at the bottom right of the A-type heatmap (light-teal). Interestingly, these junction sequences had AT-patterns in the beginning of the “left side” and the end of the “right side”. This might provide a clue to the origin of these “abnormal” junction motifs. Indeed, they may have been templated by abundant poly-T sequences in the middle of T-type 12mers flanked by alternating A-T motifs. In other words, junctions at templates of poly-A junction motifs may have been shifted by 6 nt relative to substrates. Actually, substrates have no restriction on where they may hybridize on a long template and might happen to have their ligation site in the region of poly-T of the template strand. We call this “ligation site shift”, as explained in SI section 16. Other preferred junction subsequences include repetitions of the AAT motif across the junction (the dark-teal peak in the upper left corner of the left panel).

**Fig. 4.**
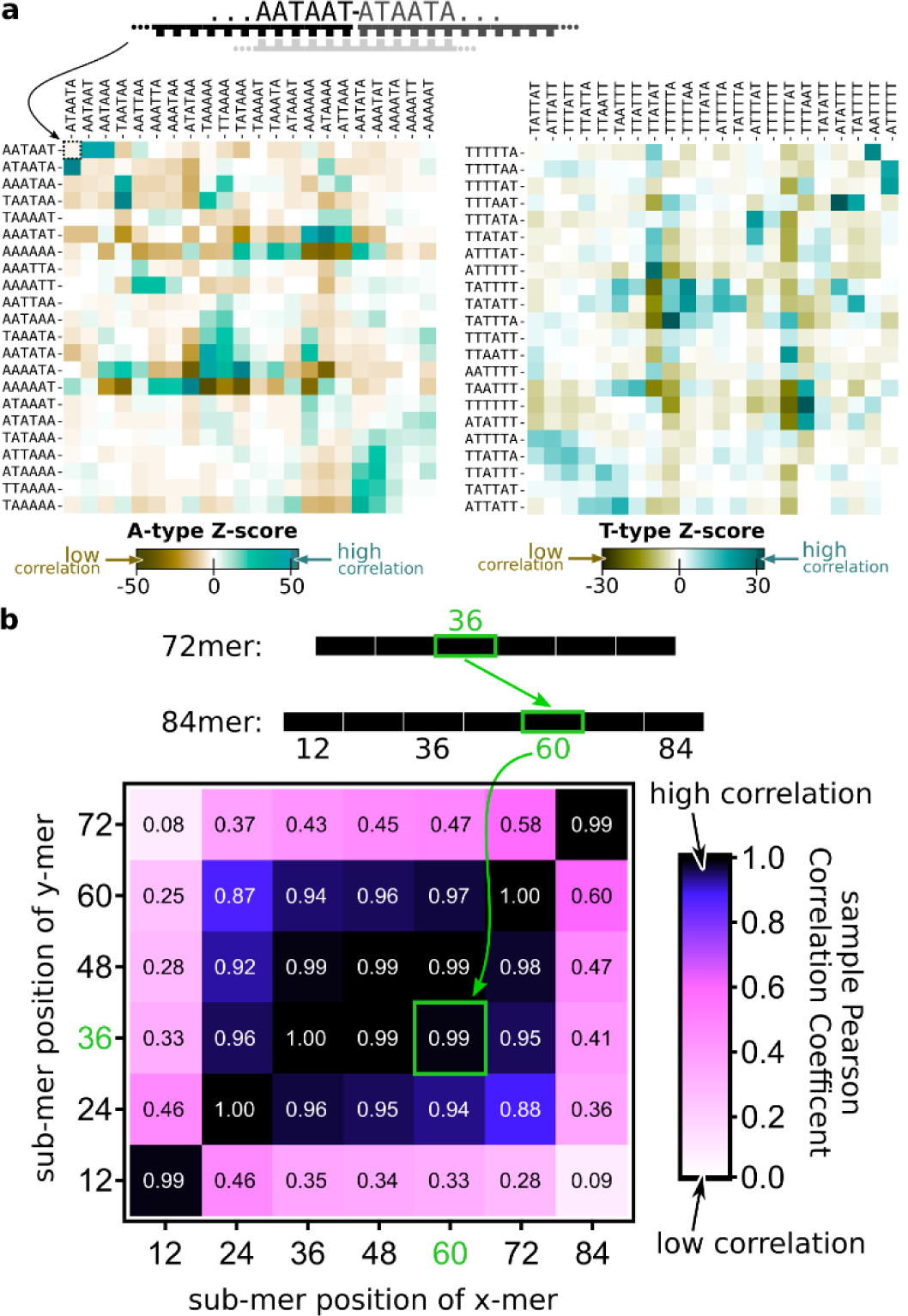
Emergent landscape of junction sequences. **a** The heatmap of Z-scores quantifying the probability to find a junction between a 6 nt sequence listed in rows followed by the 6 nt sequence listed in columns compared to finding it by pure chance and normalized by the standard deviation. Z-scores were calculated for the junction between 4^th^ to the 5^th^ 12mers in 72mers of A-type (left) and T-type (right) respectively. Other internal junctions in all long chains form very similar landscapes composed of over- (teal) and under-represented (ocher) sequences and described in detail in the text. T-type sequences complementary to A-type sequences correspond to the 90° clockwise rotation of the left panel (note a similarity of landscapes in two panels after this transformation). **b** The matrix of sample Pearson Correlation coefficients between abundances of 12mers in different positions (1 to 6) inside 72mers (rows) and 84mers (columns). Light regions mark low correlations, dark regions mark high correlations. Very high correlations (>0.9) at the center of the table mean that very similar sequences get selected at all internal positions of chains of different lengths. Different selection pressures operate on the first 12mer and the last 12mer of a chain, yet their sequences are similar in chains of different lengths.

How similar are selective pressures operating on sequences of different 12mers within longer chains? Fig. 4b quantifies this similarity in terms of sample Pearson-Correlation-Coefficient (sPCC) between abundances of 12mer sequences in different positions of long chains of different lengths. We compare the abundances of 2^12^=4096 possible 12mer sequences in positions 1 to 6 within all 72mers and compare them to each other and abundances of 12mers in positions 1 to 7 in all 84mers. Similar results were obtained for other chains longer than 36 nt. A rectangle of very high correlations (>0.9) at the center of the table in Fig. 4b means that very similar sequences get selected at all internal positions of all chains (note that only chains longer than 36nt have such internally positioned 12mers). However, the light border of the table means that a rather different subset of 12mers gets selected in the first and the last position of a multimer. Whatever the nature of selection pressure acting on these 12mers, it is consistent across oligomers of different lengths as manifested by the high correlation in the lower left and the upper right corner of the table in Fig. 4b.

A simple hypothesis comes to mind: a strand is prolonged and grows in this random sequence templated ligation system as long as the sequences attached to it share similar sequence motifs resulting in high values of sPCC for all internal 12mers. But when a 12mer sequence that is similar to the start- or end-subsequence is attached, the growth in that direction stops.

Comparison of abundances of internal 12mers in A-type and T-type subpopulations predictably yielded no positive correlation and in fact resulted in a slight negative correlation (see SI section 11). However, abundances of reverse complements of sequences from the T-type subpopulation are strongly correlated with those of the A-type resulting in a sPCC matrix similar to that shown in Fig. 4b (see SI-Fig. 12). Therefore, chains in two groups (A-type and T-type) show a considerable degree of reverse complementarity to each other. This fits the elongation and replication mechanism by templated ligation.

To further explore selection capabilities of templated ligation as a function of 12mer sequences in the initial pool we conducted three additional experiments referred to as “Replicator”, “Random” and “Network”. The “Random” experiment started with eight randomly chosen 12 nt sequences served as a control. In the “Replicator” experiment the pool consisted of eight 12 nt sequences artificially designed for efficient elongation (see below). In the “Network” experiment we populated the pool with eight naturally selected 12 nt sequences commonly found as subsequences of long strands in our original ligation experiment with 4096 12mers. To identify these 12mers, we built a network of the most common 12mers found in A-type oligomers with length of more than 48 nt. This network does not include the first and the last 12mers, in a multimer as those are known to be statistically different from the internal ones (see Fig. 4b). The circles in Fig. 5a represent unique 12 nt subsequences while their size describes their Z-scores quantifying their abundance in long chains. The width of the connecting line describes the probability that two subsequences are found one after another in a multimer. The same is done for T-type sequences (Fig. 5b). This representation of a polymer is known as de Bruijn graph^41^ and has been commonly used in DNA fragment analysis and genome assembly^42^ and more recently in the context of templated ligation^17^.

**Fig. 5.**
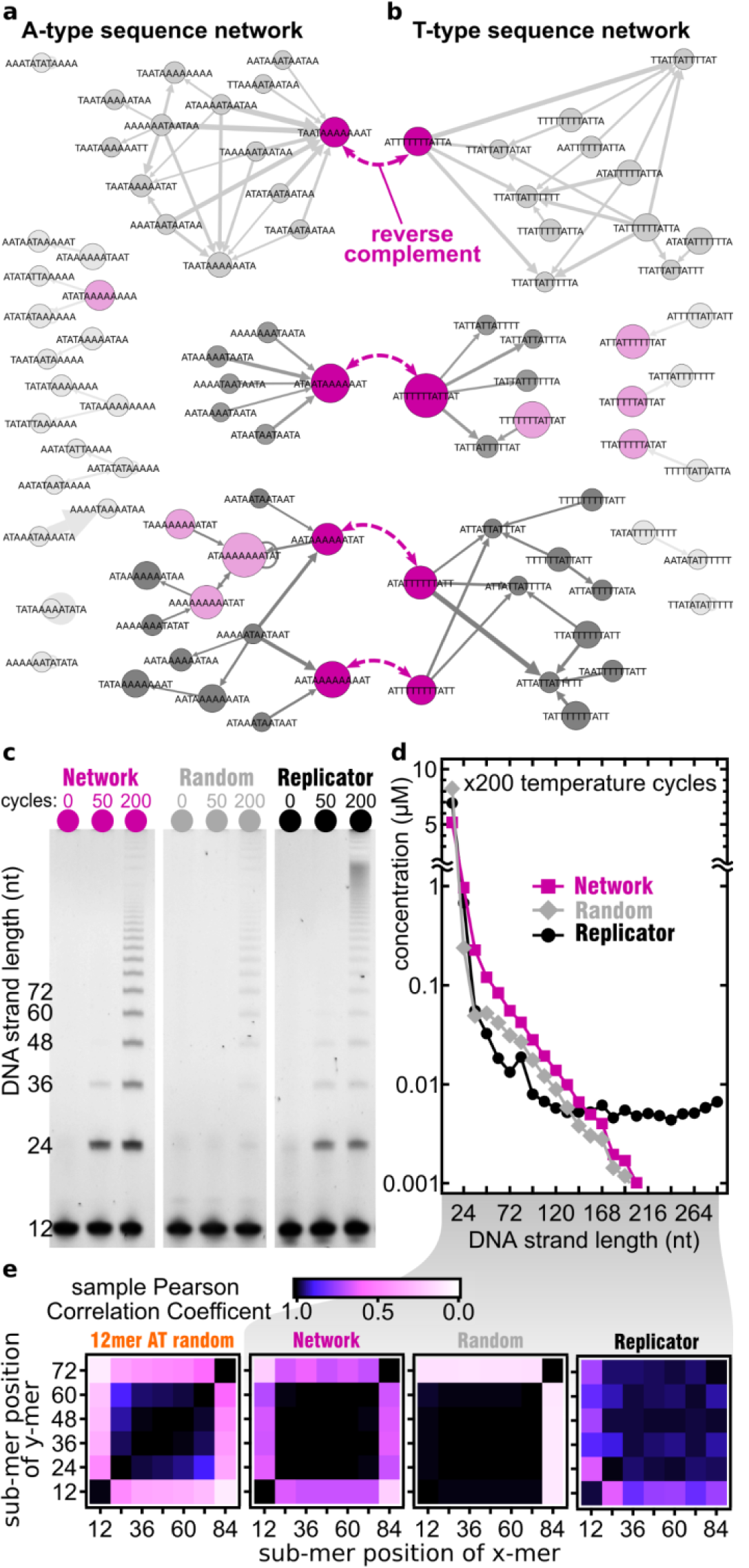
testing self-selection with custom sequence pools. **a** The de Bruijn graph of overrepresented sequence motifs between consecutive 12mers found in long oligomers. All internal junctions of A-type sequences >48 nt are shown, except the first and the last. All analyzed strands have a Z-score >30 and are sequenced at least 20 times. **b** The same de Bruijn graph but for T-type sequences with Z-score >15 and sequenced at least 10 times. Four pairs of most common reverse complementary 12mers are connected by purple dashed arrows. In each network three families with distinctly similar patterns are observed, that each include at least one of the complementary strands. Node sizes reflect relative abundance of 12mers, edge thickness denotes the Z-score of the junction between nodes it connects. Light and dark magenta-colored nodes are eight most abundant 12mers in each of two networks. **c** PAGE images of templated ligation of three different samples of 12mers after different number of temperature cycles (columns): **“Replicator”**: four substrate 12mers and four template 12mers artificially designed for templated ligation, as explained in SI, “**Random”**: eight random sequence 12mers randomly selected from the 4096 possible AT-only 12mers, “**Network”**: four most common 12mers from A-type and another four of T-type shown in dark magenta color in a). **d** After 200 temperature cycles, the “Replicator” shows a consistently higher product concentration for all lengths followed by the “Network” sample and then by the “Random” subsamples. In the “Network” and “Random” samples the length distribution above 48nt is well described by an exponential distribution as predicted in Ref ^39^. **e** Pearson correlation matrices between 12mer abundances within 72mers and 84mers in each sample (same as in Fig. 4b). While the pattern of correlations in the “Network” sample (second from left) resembles that shown in Fig. 4b (reproduced in the leftmost subpanel), the “Random” sample (second from right) singles out the last 12mer but not the first one. The “Replicator” sample (the rightmost subpanel) has its own distinct self-similar pattern of correlations.

De Bruijn networks in Fig. 5a break up into several clusters connecting 12mers with similar subsequences at junctions (TAA-TAA in the top cluster marked by a dark-magenta node, ATA-ATA in the middle one, and AAT-AAT in the bottom one). Note that these three common junction subsequences are all related via template shifts. The most common subgraphs found in the A-type network and mirrored among their reverse complements in the T-type network. This pattern is consistent with selection driven by templated ligation (see SI section 19). Among the eight most common subsequences in the A-type network (light and dark magenta nodes in Fig. 5a), four (dark magenta nodes) had a reverse complement among the eight most common subsequences of the T-type network (light and dark magenta nodes in Fig. 5b). These sequences were chosen as the pool of eight 12mers in the “Network” sample. The “Random” sample consisted of eight 12mers which were randomly chosen from the 4096 possible AT-only 12mers. The “Replicator” sample consisted of eight strands that were built to form three-strand complexes that resemble the assumed first ligation reaction in the pool (SI section 17.1).

The length distribution of oligomers (Fig. 5d) with concentrations quantified from the PAGE gel image (Fig. 5c) shows that the “Network” sample produced the most product, as the remaining 12mer sequence concentration was reduced below two other samples down to almost 5 µM. The length distribution in both “Random” and “Network” samples is well described by a piecewise-linear distribution predicted in Ref ^39^. For short product lengths ranging between 48mers up to 136mers the “Random” sample produced more oligomers than the “Replicator” sample. However, for even longer strands, the “Replicator” sample generated the largest number of really long strands since its length distribution reached a plateau around 120mers. This is probably due to the nature of the eight-sequences pools used here with the “Replicator” one made to form well aligned dsDNA that can be properly ligated. According to NUPACK^43^, 12mers in the “Random” sample should not form any complexes that could be subsequently ligated by the TAQ ligase. However, our results shown in Fig. 5c prove the existence of extensive ligation even in the “Random” sample. Presumably, it was initially triggered by small concentration of complexes formed with low probability, which were subsequently amplified due to the exponential growth of longer strands in our experiment, just like in the “Network” sample.

## DISCUSSION

We experimentally studied templated ligation in a pool of 12mers made of A and T bases with all possible sequences (2^12^=4096), subjected to multiple temperature cycles. To accelerate hypothetical spontaneous ligation reactions operating in the prebiotic world, we employed TAQ DNA ligase in our experiments. This process produced a complex and heterogeneous ensemble of oligomer products. By performing the “next generation sequencing” of these oligomers, we found that long strands in this ensemble have a significantly lower information entropy compared to a random set of oligomers of the same length. This effect became increasingly more pronounced for longer oligomers (Fig. 2e). The overall reduction in entropy was in line with the theoretical prediction obtained within a simplified model of template-based ligation^17^. In that model, the reduction of entropy was due to “mass extinction” in sequence space, with only a very limited (though still exponentially large) set of survivor sequences emerging. In the present experiment related variation in abundances of different sequences did develop but didn’t proceed all the way to extinction.

Several patterns can be easily spotted in the pool of surviving sequences. In particular, multimer strands predominantly fell in one of two groups: A-type or T-type each characterized by about 70 % of either base A or T (Fig. 2c, d). The initially single-peaked approximately binomial A:T-ratio distribution in random monomers changed into a bimodal one in longer chains. We attribute this separation into two subpopulations to the fact that such composition bias suppresses the formation of internal hairpins and other secondary structures. The self-hybridization reduces the activity of both template and substrate chains leading to a lower rate of ligation. The adaptation by separation into two subpopulations was reproduced by a kinetic model in which activities of reacting strands were corrected for hairpin formation, with realistic account for its thermodynamic cost. This model produced a bimodal distribution of A-content in 24mers, in qualitative agreement with the experimental data. Furthermore, the eventual distribution of longer oligomer lengths could be successfully captured by the maximum entropy distribution, subject to the constraint of fixed average composition of A- and T-type subpopulations. Another remarkable observation is that although formation of hairpins was suppressed through the mechanism above, a small but noticeable fraction of oligomers have extremely long stretches of internal hairpins. The likely mechanisms of their formation are either ligation of a pair of nearly complementary chains from A-type and T-type subpopulations, or self-elongation of such oligomers.

Another common pattern was a distinct AT-alternating pattern around the ligation site, as can be seen in Fig. 1b. Those AT-alternating motifs first appeared in 24-mers, and remained very common in longer chains. These features accounted for some of the reduction in sequence entropy, but did not account for all of the selection at ligation sites, where, as demonstrated by the Z-score analysis, a rich ligation landscape has developed (Fig. 4a, b). Not only some 12mers within longer chains were far more abundant than average, but there were also pairs of those that preferentially follow each other, as demonstrated by de Bruijn graphs in Fig. 5a, b.

We selected a subset of eight pairs of mutually complementary 12mers that appeared anomalously often within longer chains and were well connected within the de Bruijn graph. Using this “Network” subset as a new starting pool, we repeated the temperature-cycling experiment, and compared it to two other reference systems. One of them were eight randomly selected 12mers, the other was artificially designed to promote self-elongation. The resulting multimer population in two out of three of these pools followed a near perfect exponential length profile (Fig. 5d). The random pool resulted in a similar behavior to the network one but with significantly lower overall concentration of long chains. Both results are in an excellent agreement with theoretical predictions of reference^39^. A higher concentration of long chains generated by network 12mers indicates better overall fitness of this set compared to random 12mers. The “Replicator” set did produce a large number of very long products, presumably by a different mechanism, but a significantly smaller number of products with short and medium lengths. This indicates lower autocatalytic ability in both “Replicator” and “Random” sequence pools when compared to the “Network” pool.

For emergence of life on early earth, random oligomers needed to act in an evolution-like behavior. Here, we followed templated ligation of random 12mer strands made from two bases under temperature oscillations. Despite its minimalism, the system contains all elements necessary for Darwinian evolution: out of equilibrium conditions, transmission of sequence information from template to substrate strains, reliable reproduction of a subset of oligomer products and selection of fast growing sequences in the process. At the dawn of life, pre-Darwinian dynamics would have been important to push prebiotic systems towards lower entropy states. Such pre-selection for catalytic function could have paved the way towards eventual emergence of life.

## METHODS

### Nomenclature

***Oligomer***: a product from the templated ligation reaction with a length of a multiple of 12 nt. ***Subsequence***: 12mer long sequence in between two ligation sites or in the beginning or end of a multimer. ***Submotif***: a sequence of a certain length *x*. In contrast to a subsequence, a submotif can start at any position in a mono- or oligomers, not only at ligation sites, or the sequence start. ***Ligation site:*** in particular, the bond between two monomer or multimer strands. In context of sequence motifs, it refers to the region around this bond (±1 to 6 bases).

### Ligation by DNA ligase

For enzymatic ligation of ssDNA a TAQ DNA ligase from *New England Biolabs* was used. Chemical reaction conditions were as stated by the manufacturer: 10 µM total DNA concentration in 1x ligase buffer. The ligase has a temperature dependent activity and is not active at low (4-10 °C) and very high temperatures (85-95 °C). In our experimental system DNA hybridization characteristics are strongly temperature dependent, as shown in the SI. We expect this to have stronger influence on the overall length distribution and product concentrations than ligase activity, as the timescale of hybridization is significantly longer than the timescale of ligation (compared in SI). The manufacturer provides activity of the ligase in units/ml, specifically: “one unit is defined as the amount of enzyme required to give 50 % ligation of the 12-base pair cohesive ends of 1 μg of BstEII-digested λ DNA in a total reaction volume of 50 μl in 15 minutes at 45 °C”.

### Design of the random sequence pool

The use of a DNA ligase enables very fast ligation with low error rate. But not every DNA system is suitable for templated ligation. As stated by the manufacturer, the TAQ ligase does not ligate overhangs which are 4 nt or shorter. Therefore, the shortest possible length of strands is 10mer, opening up 4^10^ > 10^6^ different monomer sequences. The resulting pool cannot be sequenced to a reasonable extend. We artificially reduced the sequence space by limiting sequences to only include bases adenosine (A) and thymine (T). 10mer strands with random AT sequence have too low melting temperature, in a range where the ligase is not active (compare SI). We found 12mers with random AT sequences to successfully ligate and to produce longer product strands due to their elevated melting temperature. The monomer sequence space is 2^12^=4096 is not too large, so that we were able to completely sequence it multiple times.

The DNA was produced as 5’-WWWWWWWWWWWW-3’ with a 5’ POH modification by *biomers*.*net*. “W” denotes base A or T with the same probability. We analyze the “randomness” of this pool in the SI.

### Temperature Cycling

Temperature cyclers *Bio-Rad* T100, *Bio-Rad* CFX96, *Analytik Jena* qTOWER^3^ and *Thermo Fisher Scientific* ProFlex PCR System were used to apply alternating dissociation and ligation temperatures to our samples. The dissociation temperature of 75 °C was chosen, to melt short initially emerging ssDNA of up to 36mer. In the SI we also show how a variation of the dissociation temperature changes multimer product distribution in a random sequence templated ligation experiment. Lower dissociation temperatures enable us to run several thousand temperature cycles, as the stability of the TAQ DNA ligase is reduced substantially for longer times at 95 °C. Time resolution experiments with PAGE-analysis demonstrated ligase activity even after 2000 temperature cycles for a dissociation temperature of 75 °C. In experiments screening the ligation temperature (see SI), we found that for ligation temperatures of 25 °C the product length distribution is exponentially falling. For higher ligation temperatures such as 33 °C we find more long sequences, but almost no 24mer and 36mer sequences. For sequenced samples we chose a ligation temperature of 25 °C because the library preparation kit is better suited for shorter DNA strands. In sequencing data for samples with 33 °C the yield was very low, but the results are similar to the sequencing data of samples with 25 °C ligation temperature, but with comparably worse statistics. For dsDNA dissociation in each temperature cycle the corresponding temperature is held for 20 s with subsequent 120 s at the ligation temperature.

### Sequencing by Next Generation Sequencing (NGS)

For sequencing we used the Accel-NGS 1S Plus DNA Library Kit from *Swift Biosciences*. The sequencing was done using a HiSeq 2500 DNA sequencer from *Illumina*. The kit was used as stated in the manufacturer’s manual. All volumes were divided by four to achieve more output from a limited supply of chemicals. Library preparation was done in four steps: first a random sequence CT-tail was added to the 3’ end of the DNA by (probably, the manufacturer does not give information about this step) a terminal transferase. In a single 15 min ligation step the back primer sequence (starting with AGAT…) was ligated to the 3’ end of the random CT-stretch. In the second step a single cycle PCR was used to produce the reverse complement and to leave double stranded DNA with a single A overhang. Step three ligated the start primer to the 5’ end of the DNA. Step four added barcode indices to both ends of the DNA by a PCR reaction. This step was done several times to result in the desired amount of DNA for sequencing.

### Sequence Analysis

Demultiplexing was done by a standard demultiplexing algorithm on servers of the Gen Center Munich running an instance of Galaxy^44^ connected to the sequencing machine. *Illumina*-sequencing creates three FASTA-files, listing the front and the back barcodes and the read sequence, for each lane of the flow cell. The demultiplexing-algorithm matches the barcodes of the prepared library DNA to the read sequence and produces a single FASTA file including the read quality scores.

The sequence-data was analyzed with a custom written *LabVIEW* software. The main challenge was to separate the read sequences from the attached primers. The start primer is automatically cut in the demultiplexing step. The end primer is cut with an algorithm based on regular expression (RegEx) pattern matching. With RegEx we first search for multiples of the monomer length. If these structures were followed by at least four bases of C or T followed by the sequence AGAT we concluded that we found a relevant sequence. The 3’-primer was cut and the resulting sequence saved for analysis.

RegEx for searching AT random sequences:

(^[ATCG]{12}|[ATCG]{24}|[ATCG]{36}|[ATCG]{48}|[ATCG]{60}|[ATCG]{72}|[ATCG]{84})(?=([CT]{4,}AGAT))

RegEx for selecting a maximum of X false reads of G or C in random sequence AT samples: ^(?!(?:.*?(G|C)){X,})^([ATCG]{12,}). The sequenced library may have primer-primer dimers and oligomers as well as partial primers that were falsely built in the library preparation step. As the SWIFT kit is made for longer sequences by design, shorter sequences such as 12mer in our study may have lower yields and larger error rates for the library kit chemistry. Therefore, the inclusion of sequences with a single or multiple false reads can improve the statistics, as long as submotifs with obviously faulty reads are ignored in the analysis.

## Author’s contribution

P.W.K. performed the experiments, prepared the library for sequencing, performed the demultiplexing, the analysis, programmed the analysis software, analyzed the data, drafted and wrote the manuscript. A.V.T and S.M. performed the theoretical analysis and analyzed the data in context of their already published theoretical work, drafted graphs, drafted and wrote the manuscript. D.B. contrived the experiment, guided the experimental progress, analyzed data and drafted the manuscript.

## Acknowledgements

The authors would like to acknowledge funding by the Deutsche Forschungsgemeinschaft (DFG, German Research Foundation)– Project-ID 201269156 – SFB 1032, the Advanced Grant (EvoTrap #787356) PE3, ERC-2017-ADG from the European Research Council, CRC 235 Emergence of Life (Project-ID 364653263) and the Center for NanoScience (CeNS). We would like to thank Ulrich Gerland, Tobias Göppel, Joachim Rosenberger and Bernhard Altaner for their helpful remarks and discussions about hybridization energies, baseline corrections and interpretation of multimer product distributions. P.W.K and D.B. thank Stefan Krebs and Marlis Fischalek at the Gene Center Munich for help with the library preparation and the sequencing the samples and Annalena Salditt and Filiz Civril for comments on the manuscript. This research was partially done at, and used resources of the Center for Functional Nanomaterials, which is a U.S. DOE Office of Science Facility, at Brookhaven National Laboratory under Contract No.∼DE-SC0012704.

